# PathFold: Predicting the Entire Protein Folding Pathway from Protein Sequence Alone

**DOI:** 10.64898/2026.08.26.747321

**Authors:** Zicong Zhang, Nabil Ibtehaz, Yuki Kagaya, Ziyi Xu, Pranav Punuru, Daisuke Kihara

## Abstract

Recent advances in protein structure prediction, exemplified by AlphaFold, have largely addressed the determination of static structures, one aspect of the protein folding problem. However, predicting folding pathways, by which proteins reach their native states, remains a significant challenge. Here, we present PathFold, a deep learning framework that predicts protein folding pathways directly from sequence information. PathFold leverages an AlphaFold-based module to extract structural information from the sequence and generates a progressive folding trajectory from an extended conformation using a diffusion model. By modeling the full trajectory, it enables prediction of folding intermediates and transition pathways, analogous to those observed in steered molecular dynamics (SMD) simulations. The predicted pathways reveal well-defined intermediates and sequential folding events, and show agreement with experimental folding data, including measured Φ-values.

## Introduction

Recent success in deep learning–based protein structure prediction^1^ have enabled highly accurate prediction of fully folded three-dimensional structure of proteins from amino acid sequences, substantially advancing the static structural component of the protein folding problem^2^. However, these methods do not explain how proteins fold into their native structures from unfolded conformations^3^. Consequently, predicting protein folding remains a major challenge; the heart of the protein folding problem, the folding process component of the protein folding problem that has been studied in biophysics for decades continues to pose significant difficulty^4,5^.

The protein folding has been extensively studied over the past decades using a wide range of experimental approaches that characterizing folding pathways and intermediate states, including rapid-mixing methods^6^, hydrogen-exchange experiments^7^, single-molecule FRET^8^, optical tweezers^9^, ultrafast temperature-jump spectroscopy^10^, and small-angle X-ray scattering (SAXS) studies^11^. In the so called Φ-value analysis the effects of point mutations on folding kinetics and stability are used to infer the extent of native-like structure formed in the folding transition state^12^. Computationally, molecular dynamics (MD) simulations have been used to investigate atomic details and thermodynamics of the folding process^13,14^. In addition to regular MD simulations, steered molecular dynamics (SMD) simulations, which apply external forces to pull protein chains, have been widely used to investigate a range of biophysical processes, including protein folding and unfolding mechanisms^15,16^. SMD can simulate atomic force microscopy (AFM) pulling experiments, which measure the forces generated during the pulling of a protein chain^17,18^. Advantages of SMD include its ability to be directly compared with single-molecule AFM experimental data, as well as its accelerated sampling of conformational changes, which provides residue-level mechanistic insight into structural transitions. SMD has been also been successfully applied to investigate protein unbinding^19^, heterogeneous unfolding pathways^20^, and peptide–receptor interactions^21^, establishing SMD as a reliable approach for modeling AFM-based mechanical measurements at atomic resolution.

In the past few years of the post-AlphaFold era, deep learning has been applied to many modeling and prediction tasks related to proteins and other biomolecular structures^22–28^ and design^29–32^. Among these tasks, a major focus has been modeling protein flexibility at various scales. AlphaFlow^33^ combines AlphaFold with flow-based generative modeling to learn and sample sequence-conditioned protein conformational ensembles beyond single static structures. BioEMU employs deep generative models to reproduce equilibrium conformational ensembles learned from molecular dynamics simulations^34^. aSAM is a latent diffusion–based generative model trained on molecular dynamics data, with a temperature-conditioned variant that explicitly captures temperature-dependent protein conformational ensembles^35^. Although these methods have been successful in modeling conformational ensembles, they are not designed to study protein folding and do not generate ordered folding trajectories.

Here, we present PathFold, the first deep learning framework for protein folding pathway prediction. PathFold operates at the Cα level and employs a diffusion-based architecture conditioned on AlphaFold-derived single and pair representations, which provide information about the native structure that serves as the endpoint of folding. Given a protein sequence and a fixed number of preceding intermediate states, PathFold predicts the next conformation and iteratively generates an ordered trajectory from a fully extended chain toward the native structure. A fundamental challenge in developing such a model is the lack of large-scale, atomistic protein folding trajectories for training: experimentally, complete folding pathways cannot be directly observed at atomic resolution, while computational generation of sufficiently long folding trajectories remains prohibitively expensive for large-scale dataset construction. To circumvent this data bottleneck and establish the feasibility of learning folding pathways, we generated steered molecular dynamics (SMD) trajectories in which proteins are mechanically unfolded and reversed these trajectories to provide supervised folding pathways. Although these reversed trajectories do not represent genuine folding dynamics, they provide a practical surrogate for developing and testing a framework for trajectory learning in the absence of large-scale folding trajectory datasets.

Evaluating folding pathway predictions requires metrics beyond static structure comparison, as conventional measures of structure comparison such as root mean square deviation (RMSD) are insufficient for assessing sequential folding progression and intermediate-state alignment. To address this limitation, we introduce a trajectory alignment scoring metric that compares predicted trajectories with reference trajectories derived from SMD simulations by aligning intermediate states and quantifying folding progress through native contact formation.

We evaluated PathFold on a benchmark of 175 proteins with sequence lengths ranging from 29 to 399 residues and SMD trajectories containing 62 to 1,612 frames. Using the trajectory alignment scoring metric developed in this study, PathFold consistently generated ordered folding pathways with high trajectory alignment scores. In addition, PathFold produced intermediate protein states with high similarity to those observed in SMD simulations and consistently predicted forward folding trajectories. We further examined three well-characterized proteins, WW domain^36^, cold shock protein^37^, and Ubiquitin^38^, whose folding mechanisms have been extensively studied experimentally. Residue-level folding scores derived from the predicted trajectories showed strong correspondence with experimental Φ-value profiles, demonstrating that PathFold captures key features of folding transition states.

## Results

### Overview of PathFold Architecture

**Figure 1** illustrates the overall workflow of PathFold, including the inference loop for folding trajectory prediction (**Fig. 1a, b**) and the training paradigm (**Fig. 1c**). PathFold takes a target protein sequence together with a starting fully extended conformation and generates a protein folding trajectory, which consists of an ordered series of protein structure frames leading to the native structure. From the input sequence, single and pair representations are first computed using the AF2 Evoformer^1^. PathFold builds upon the Genie^39^ framework, a diffusion-based protein design model originally developed for backbone generation. While Genie focuses on de novo backbone design, PathFold extended its architecture to incorporate additional conditioning signals, including AF2 single and pair representations as well as intermediate structural states (**Supplementary Algorithm 1**). These modifications enable PathFold to generate protein-specific folding pathways rather than generic backbone structures. The detailed architecture of PathFold is presented in **Supplementary Fig. S1**.

**Fig. 1.**
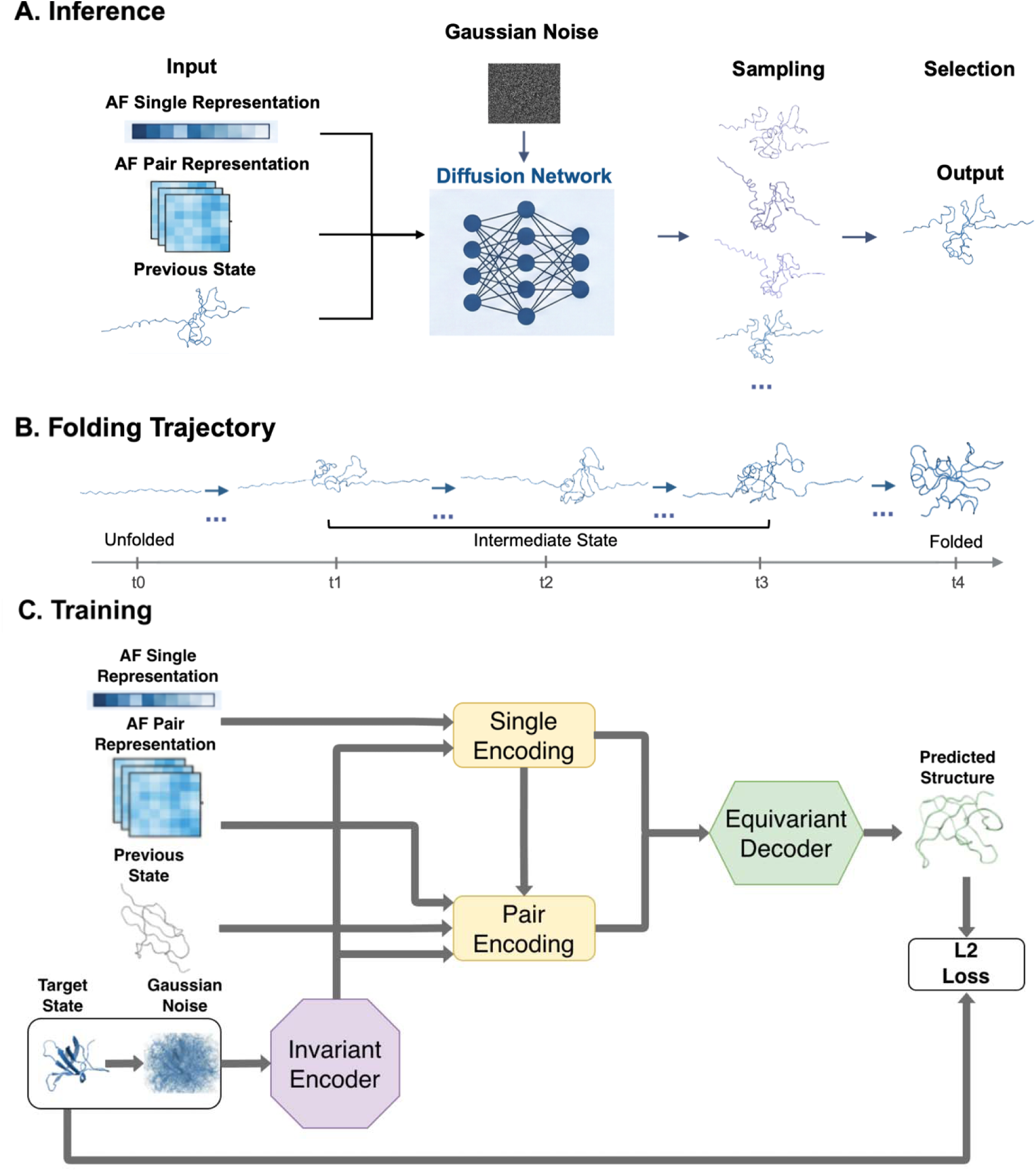
Inference and training frameworks of PathFold. **a**, the overview of the inference pipeline. The PathFold model takes as input (1) AF2 single and pair representations derived from the target protein sequence, and (2) the current intermediate state. PathFold employs a diffusion network to generate candidate intermediate states that progressively fold the protein. The final structure is selected from the candidate pool based on predefined criteria, which considers contact-map progression toward the folded state while preventing abrupt structural transitions. (see Methods). The selected intermediate state is then fed back into the model for subsequent iterations. **b**, PathFold generates a folding trajectory starting from a fully extended state and iteratively predicts successive intermediate states until the protein reaches its native folded structure. A generated trajectory for PDB ID: 1A2P is shown as an example. **c**, Overview of the PathFold training procedure. In addition to the single and pair representations generated by the AF2 Evoformer, we used two Cα structures from SMD simulations separated by 50 frames. The more folded conformation was used as the target state and was perturbed with Gaussian noise during diffusion training, whereas the less folded conformation serves as the previous state. The AF2 representations and the previous-state structure were encoded and fused with single and pair encodings produced by an invariant encoder, which processes the current noisy structure sampled during the diffusion process around the target state. An equivariant decoder then integrates the single and pair encodings to predict the denoised target structure. The predicted structure is supervised using a per residue L2 distances loss against the ground-truth target state.

We train PathFold using a dataset of (un)folding simulations using SMD, which contains 940 proteins spanning a wide range of sequence lengths from 29 to 399 residues (Methods). The SMD simulations started from the native structure of the target proteins, with one terminus restrained while the other was pulled using a harmonic spring^40^. The simulations proceeded until the protein reached a fully extended conformation in which each residue primarily interacted only with its immediate sequence neighbors.

To construct the training dataset, each SMD trajectory is decomposed into individual time frames, with each frame representing an intermediate structural state along the unfolding pathway (**Fig. 1c**). During training, we sample pairs of structures from the same SMD trajectory that are separated by 50 frames. The more folded structure is used as the target state, while the less folded structure serves as the previous state, whose contact map is encoded as input to the model. The target state is encoded using an SE(3)-invariant encoder based on the Frenet–Serret (FS) frame^41^, which captures local geometric features while preserving rotational and translational invariance, together with diffusion-step information. This encoder generates single encodings, representing residue positions along the protein sequence, and pair encodings, capturing pairwise distances derived from Cα coordinates. These encodings are combined with projected AF2 single and pair representations, which provide additional input information to the network. On the other hand, the previous state is represented with a residue contact map and fused into the pair encodings to provide structural context for the folding progression. The fused single and pair representations are then passed to an SE(3)-equivariant decoder that employs Invariant Point Attention (IPA), similar to the structure module used in AlphaFold2 (AF2). The decoder predicts the Cα coordinates of the next intermediate folding state, which are compared with the ground-truth target state using per residue L2 distances loss. Because many different folded conformations were used as target states during training, the model effectively learns the general progression of protein structure formation during folding.

We trained PathFold on a dataset of 940 proteins comprising 728,597 SMD simulation frames (**Supplementary Table S1**) and validated it on an independent set of 175 proteins with 69,953 SMD simulation frames (**Supplementary Table S2**). The training and validation sets were split to ensure low sequence redundancy, with no pair sharing more than 25% sequence identity. For benchmarking, we evaluated PathFold on 175 proteins containing 99,172 SMD simulation frames (**Supplementary Table S3**). The number of frames in the SMD frames is proportional to the protein length (**Supplementary Fig. S2)**. Additional details on dataset construction are provided in the Methods section.

For inference, PathFold starts from a fully extended protein conformation together with AF2 single and pair representations. The folding trajectory is produced by iteratively predicting the next intermediate structural state, which is then used as the previous state for the next iteration (**Supplementary Algorithm 2)**. At each iteration, the diffusion model simultaneously generates multiple candidates of the next-state structure in a single forward sampling pass using a configurable batch size (set to 10). These sampled structures form a candidate pool from which a single structure is selected using a folding progression–based model selection strategy (Methods). Folding progression is defined as the change in contact-map similarity between the candidate structure and the folded structure relative to the previous iteration. To encourage steady folding progression and prevent abrupt structural jumps, at each iteration the structure with the largest progression under a predefined threshold is selected as the next intermediate state. In addition to the base model which takes a single previous state as input, which is terms as the PathFold-1 model, we further developed two more PathFold variants that incorporate three or six preceding intermediate structures (PathFold-3 and PathFold-6 models). To generate folding trajectory movies, predicted intermediate structures were first checked for mirror inversion relative to the AF2-predicted folded structure using a backbone handedness criterion based on local Cα cross products (**Supplementary Algorithm 3)**. Mirrored structures were corrected by reflecting the coordinate system. Consecutive structures were then connected using the PyMOL^42^ morph function, producing interpolated multi-state trajectories for movie rendering (**Supplementary Algorithm 4)**. Inference is computationally efficient, with PathFold generating folding trajectories for proteins of up to 400 residues in under 45 min (**Supplementary Fig. S3**).

### Predicted trajectories by PathFold

We first illustrate representative folding trajectories generated by PathFold (**Fig. 2**). Predicted trajectories were aligned to the corresponding SMD trajectories using PathScorer, a trajectory alignment method developed in this study (Methods). PathScorer performs global trajectory alignment using dynamic programming based on pairwise structural similarity scores between frames. Structural similarity was quantified using contact maps, yielding scores between 0 and 1, with 1 indicating identical contact maps.

**Fig. 2.**
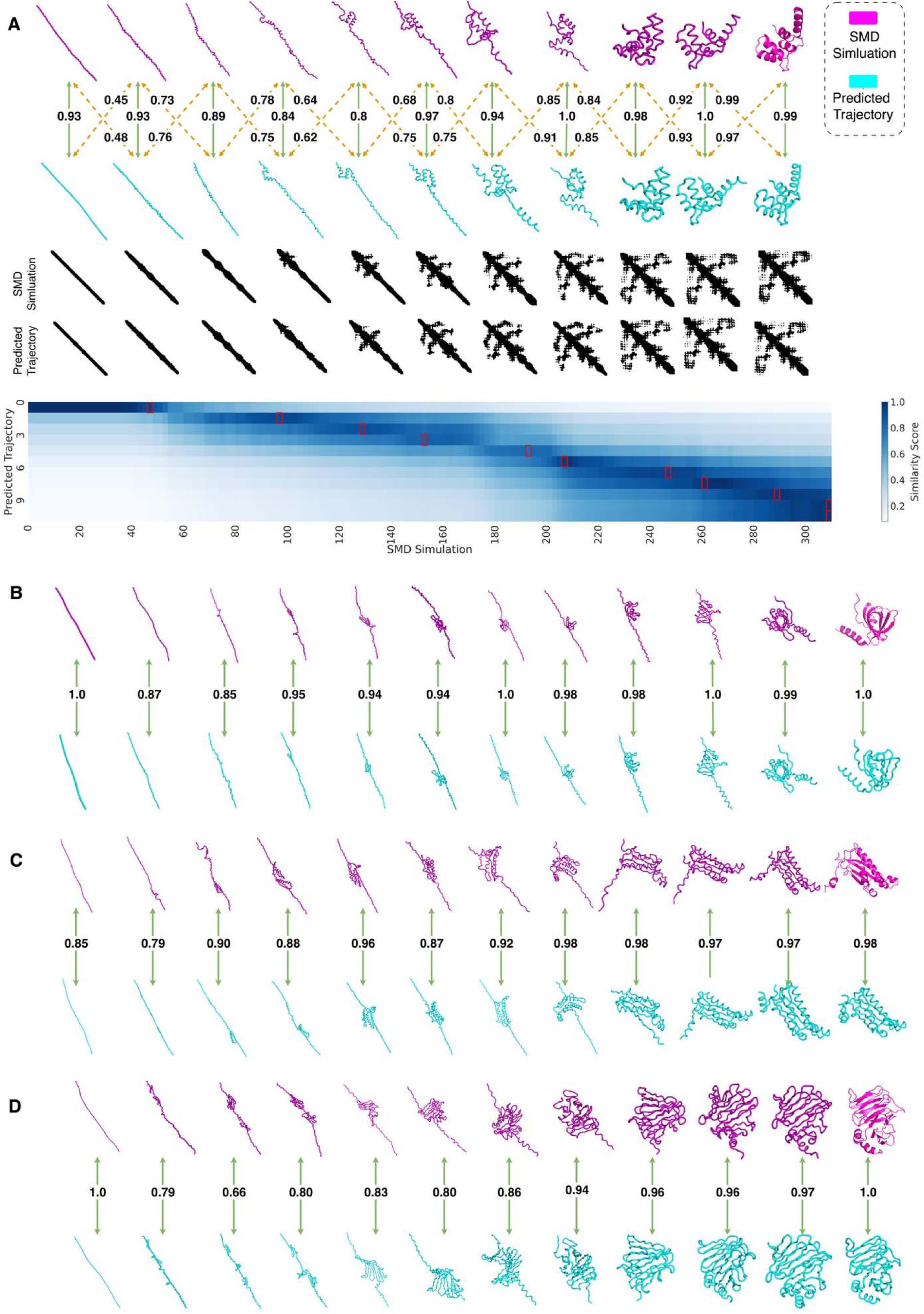
Example of predicted protein folding trajectories and alignment against the corresponding SMD trajectories. Comparison of SMD trajectories (magenta) with predicted trajectory by PathFold (cyan). Each column denotes successive time steps from a stretched conformation to the folded structure. Solid green arrows indicate pairs of frames between two trajectories that are matched by PathScorer and dashed yellow arrows show some pairs of neighboring frames. Numbers are similarity scores based on their contact maps. **A**, calmodulin-like domain of CsTAL3 (5X2E_A), 80 residues. The SMD trajectory contains 311 frames. The middle rows, residue–residue contact maps of the SMD and the predicted trajectories at corresponding stages. The bottom heatmap reports pairwise similarity scores between SMD frames (x-axis) and predicted trajectory states (y-axis). Darker blue means alignment scores closer to 1 and light blue means alignment score closer to 0. Red boxes indicate the highest-similarity SMD frame for each predicted trajectory state. **B**, CST complex subunit TEN1 (4JOI_D), 116 residues. 463 SMD frames. **C**, 5-carboxymethyl-2-hydroxymuconate delta-isomerase, (4JJ9_B), 151 residues. 600 SMD frames. **D**, Membrane protein insertase YidC (5Y82_C), 199 residues. 799 SMD frames. Contact maps and alignment heat maps for panel B-D are provided in **Supplementary Figure S6**.

To contextualize the alignment score, we generated five independent SMD trajectories for each test protein and computed pairwise alignment scores among them. Pairwise trajectory comparisons are shown in **Supplementary Fig. S4**, and the score distribution is summarized in **Supplementary Fig. S5**. Independent SMD simulations do not yield identical trajectories but they exhibited a high average alignment score of 0.894. Predicted trajectories achieved slightly lower but comparable scores, with average alignment scores of 0.804, 0.815, and 0.733 for PathFold models using one, three, and six previous frames, respectively.

In **Fig. 2**, we show four examples of predicted folding trajectories. The first example (**Fig. 2A**) is calmodulin-like domain of CsTAL3 (PDB ID: 5X2E_A) with 80 amino acid residues. This structure consists of five short α-helices, residue 1–7 (helix 1), 19–26 (helix 2), 36–45 (helix 3), 51–58 (helix 4), and 66–76 (helix 5). The folding in SMD simulation (magenta) starts with helix 2 and 3 and the event continues to include helix 1 region although it is not fully formed yet at this point. Helix 4 is formed subsequently and the trajectory ends with simultaneous formation of helix 1 and folding helix 5. The five SMD trajectories are consistent with a high average path similarity score of 0.925. In this example, the predicted folding trajectory (cyan) closely follows the SMD trajectory. Similarity scores are shown between corresponding frames from SMD and predicted trajectories (green arrows) as well as neighboring frames (yellow dashed lines). Corresponding frames have higher scores than between neighboring frames showing unambiguous folding progress of the predicted trajectory, which closely agree with SMD. The middle row shows the corresponding contact maps of the frames, which provides additional view of evolving residue-residue interactions in the folding process. Pairwise scores of frames from the SMD (x-axis) and the PathFold’s trajectories (y-axis) are visualized in the third row. The red boxes show frames that are aligned by PathScorer alignment algorithm. Overall, the predicted trajectory captured the key folding events with a close agreement with SMD, achieving a high alignment score of 0.93.

The next example (**Fig. 2B and Supplementary Movie 1**) is the human telomeric CST complex subunit TEN1 (4JOI_D; 116 residues), which has a solenoidal β-sheet with five β-strands at residues 25–96 (β1, res. 30–36; β2, res. 38–48; β3, res. 50–56; β4, res. 71–77; β5, res. 80–90) and a C-terminal α-helix (residues 106–116). The folding of this protein is as sequential formation of the complete solenoidal β-sheet region, followed by folding of the helix. The formation order within the solenoidal β-sheet region varied across the five MD trajectories: three trajectories formed β1–β2 interaction first before β3–β5, whereas other two trajectories started from β3–β5 formation before β1–β2. The predicted trajectory by PathFold captured the β1–β2-first pathway. Out of 12 frames predicted by PathFold, β1-β2 interaction was formed at frame 4 with a high similarity score of 0.95 to the corresponding SMD frame. The full solenoidal β-sheet was folded by the frame 7 with a very high similarity of 0.995 to the corresponding SMD frame. Finally, the helix was formed in frames 8–12.

**Fig. 2C** is a folding path of an α/β class protein, 5-carboxymethyl-2-hydroxymuconate delta-isomerase (PDB ID: 4JJ9_B) with 151 residues. The SMD trajectory has 600 frames. As an α/β class protein, it has an alternating arrangement of secondary structures; β-strand 1 at residues 6–16), α-helix 1 (res.26–38), β-strand 2 (res. 45–56), β-strand 3 (res. 81–91), α-helix 2 (res. 94–118), and β-strand 4 (res. 121–131). For this protein, all five SMD trajectories exhibit a highly consistent folding pattern with an average path score of 0.903. Starting from the extended state, the first major folding event occurs rapidly between SMD frames 210 and 240, characterized by the formation of α-helix 2 together with β-strands 3 and 4, which assemble into an initial folding unit. This folding intermediate is subsequently expanded through incorporation of α-helix 1 and β-strand 2 during frame 390–435. Finally, β-strand 1 folds and packs against the existing structure, completing the native β-sheet architecture between the SMD frame 480 and 600. PathFold successfully captures this folding pathway. PathFold identified the initial formation of α-helix 2 and β-strands 3–4, corresponding to SMD frame 421 with a contact similarity of 0.877. The subsequent incorporation of α-helix 1 and β-strand 2 was captured by PathFold at time frame of 6 to 8, matching SMD frames 478, 491, and 516 with contact similarities of 0.874, 0.915, and 0.979, respectively. Finally, the incorporation of β-strand 1 into the preformed β-sheet was captured at times frame 9, corresponding to SMD frame 591 with a contact similarity of 0.966.

The final example (**Fig. 2D**) highlights a case where PathFold predictions did not fully agree with the SMD trajectories. The target, membrane protein insertase YidC (PDB ID: 5Y82_C; 199 residues), folds within 799 SMD frames and adopts a β-supersandwich composed of 15 β-5strands (res. 1–171) followed by two C-terminal α-helices (res. 175–181 and 186–196). For analysis, we divided the β-supersandwich into three regions: region 1 (res. 12–48), region 2 (56–96), and region 3 (116–150). In all five SMD trajectories, the β-supersandwich forms before the C-terminal helices. However, the order of early folding events varies: in three trajectories, region 2 forms first and nucleates assembly of the β-supersandwich, whereas in the remaining two, region 1 forms first. Despite this variability, all trajectories converge on a common late-stage event in which region 3 first forms a local β-sheet and subsequently integrates into the folding unit formed by regions 1 and 2 between frames 568 and 643.

PathFold predicted a folding pathway that was not observed in the SMD trajectories. Specifically, regions 1–3 formed independently at an early stage (frame 3), after which regions 2 and 3 associated to form an intermediate folding unit that subsequently incorporated region 1 to complete the β-supersandwich. Consequently, agreement with the SMD trajectories was limited during the early folding stage; PathFold frame 3 achieved a maximum contact similarity of 0.710 to SMD frame 582. Once the β-supersandwich was assembled, however, PathFold closely recapitulated the remaining folding events. In particular, formation of the two C-terminal α-helices at PathFold frames 7–9 matched SMD frames 758, 780, and 785 with contact similarities of 0.964, 0.974, and 0.961, respectively.

### Trajectories of Multi-Domain Proteins

Next, we show examples of folding trajectories of multi-domain proteins (**Fig. 3**). The first example is the two-domain Protection of Telomeres 1 (POT1) protein (PDB ID: 5UN7_A; Fig. 3A and Supplementary Movie 2), which consists of 302 residues and folds over 1,257 SMD frames. This protein contains an oligonucleotide/oligosaccharide-binding (OB) fold domain (residues 1–60 and 208–302) and a Holliday Junction Resolvase (HJR) domain (residues 61–207), with the HJR domain inserted into the OB-fold domain^43^. All five SMD trajectories are highly consistent (average trajectory similarity score: 0.896). Analysis of the SMD trajectories reveals a hierarchical folding process. The HJR domain folds first, reaching its native-like conformation between SMD frames 282 and 735. This is followed by the folding of the second segment of the OB-fold domain (res. 208–302) between frames 800 and 1127, while the first segment of the OB-fold domain (res. 1–60) folds over frames 761 to 1257. Together, these observations indicate that formation of the HJR domain precedes assembly of the complete OB-fold domain. PathFold successfully captured the major folding events along this pathway. At frame 4, the formation of the HJR domain was identified with a structural similarity of 0.849 to SMD frame 814. Subsequent folding of the OB-fold domain was captured at frames 6, 7, and 8, with similarities of 0.953, 0.934, and 0.924 to SMD frames 1037, 1143, and 1194, respectively. Fig. 4A visualizes domain-specific similarity of predicted folding intermediates to the native structure. Consistent with the SMD trajectories, the HJR domain achieves a high similarity score at earlier frames, whereas the OB-fold domain reaches its native conformation only at later stages.

**Figure 3.**
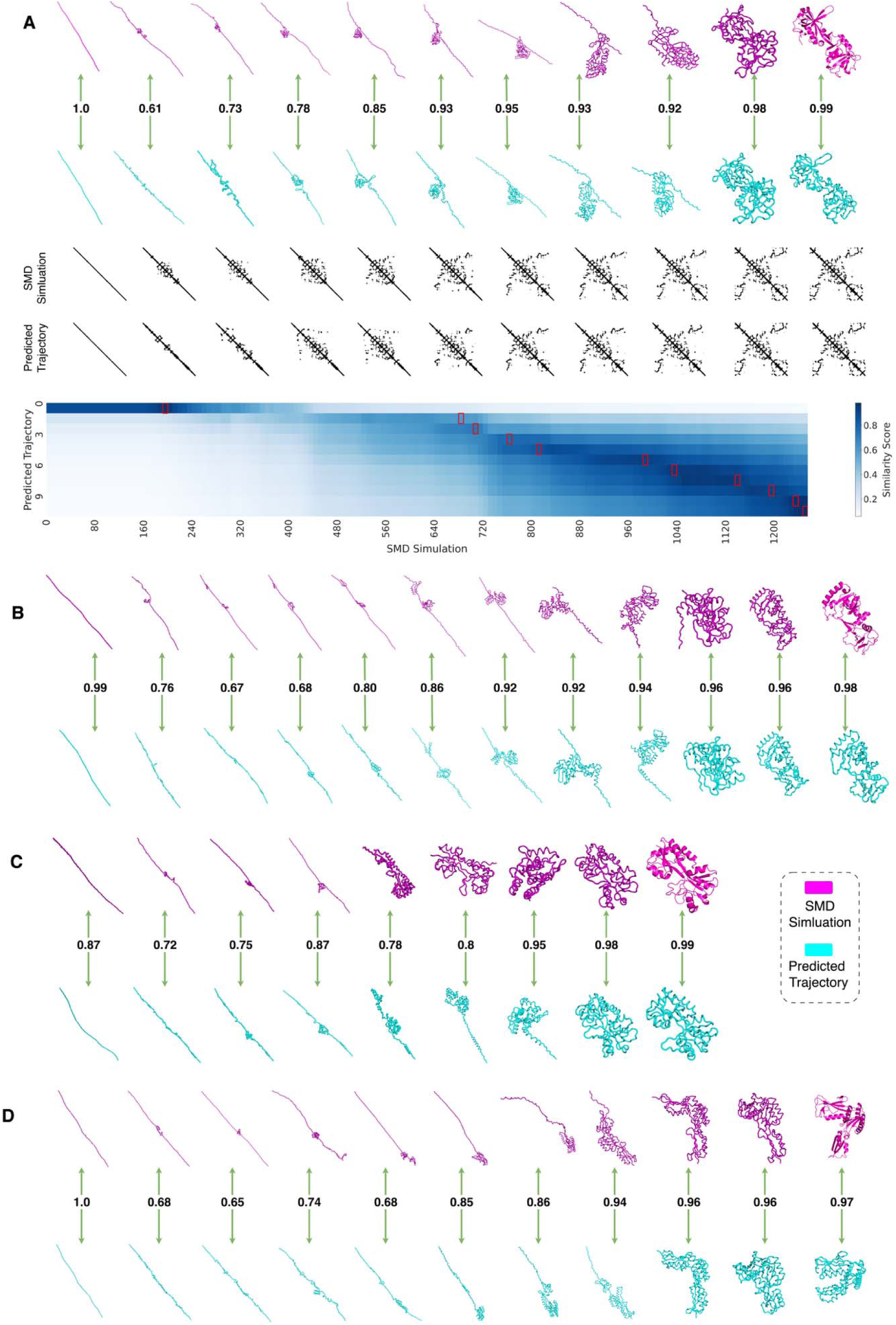
Examples of trajectories of multi-domain proteins. Comparison of SMD trajectories (magenta) and the PathFold-predicted trajectory (cyan). Each column represents a successive time step from the stretched conformation to the folded structure. Solid green arrows connect pairs of matched frames identified by PathScorer, with numbers indicating their contact-map similarity scores. For each protein, the SMD trajectory with the highest alignment score among the five generated trajectories is shown. **A**, two-domain Protection of Telomeres 1 (POT1) protein (PDB ID: 5UN7_A), 302 residues and 1257 frames of MD trajectory 1 (blue in Fig. 4A). The middle rows, residue–residue contact maps of the SMD and the predicted trajectories at each corresponding stage. The bottom heatmap reports pairwise similarity scores between SMD frames (x-axis) and predicted trajectory states (y-axis). Darker blue means alignment scores closer to 1 and light blue means alignment score closer to 0. Red boxes indicate the highest-similarity SMD frame for each predicted trajectory state. **B**, Siderophore Biosynthesis Protein SbnI (PDB ID: 5UJD_B), 254 residues. 1035 frames of MD2 (red in Fig. 4B). **C**, Extracellular solute-binding protein (PDB ID: 5SV6_A), 246 residues. 995 frames of MD5 (orange in Fig. 4C). **D**, *Arabidopsis thaliana* Toc75 POTRA domains (PDB ID: 5UAY_A), 302 residues. 1230 frames of MD1 (blue in Fig. 4D). Contact maps and alignment heat maps for panel B-D are provided in **Supplementary Figure S7**.

**Figure 4.**
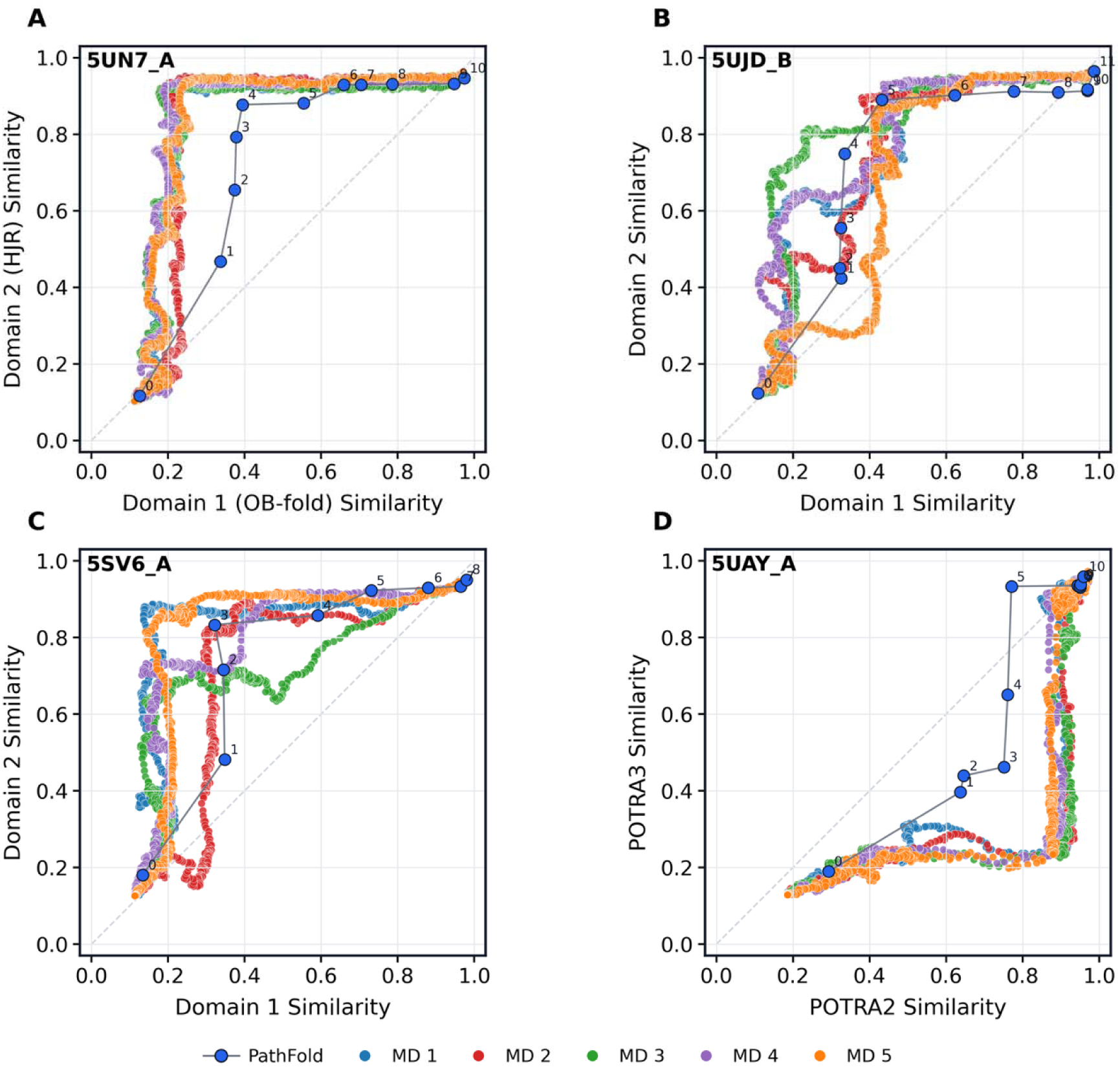
Comparison of MD and PathFold folding trajectories using domain-level structural similarity. The PathFold trajectory is shown as blue circles connected by a line, where frame 0 represents the initial extended state and frame 10 represents the folded structure. Five MD trajectories are shown for comparison: MD1 (blue), MD2 (red), MD3 (green), MD4 (purple), and MD5 (orange). Domain similarity was calculated as the Dice similarity between the contact map of a domain in a PathFold-predicted structure or MD frame and the corresponding contact map in the native folded structure. **A**, Protection of Telomeres 1 (POT1) protein (PDB ID: 5UN7_A), a two-domain protein containing an oligonucleotide/oligosaccharide-binding (OB) fold domain (residues 1–60 and 208–302) and a Holliday Junction Resolvase (HJR) domain (residues 61–207). The OB-fold similarity is shown on the x-axis and the HJR similarity on the y-axis. **B**, Siderophore biosynthesis protein SbnI (PDB ID: 5UJD_B), which contains Domain 1 (residues 1–83 and 205–240), Domain 2 (residues 84–204), and Domain 3 (residues 241–254). The similarity of Domain 1 is shown on the x-axis and the similarity of Domain 2 on the y-axis. **C**, Extracellular solute-binding protein (PDB ID: 5SV6_A). This protein consists of two domains: Domain 1 (D1; residues 1–77 and 195–246) and Domain 2 (D2; residues 88–194). The similarity of Domain 1 is shown on the x-axis and the similarity of Domain 2 on the y-axis. D. Arabidopsis thaliana Toc75 POTRA domains (PDB ID: 5UAY_A). This protein contains three tandem polypeptide transport-associated (POTRA) domains: POTRA1 (residues 34–103), POTRA2 (residues 104–193), and POTRA3 (residues 194–302). The similarity of POTRA2 is shown on the x-axis and the similarity of POTRA3 on the y-axis.

The second example is a three-domain structure, Siderophore Biosynthesis Protein SbnI (PDB ID: 5UJD_B; Fig. 3B and Supplementary Movie 3) ^43^. It is 254 residue-long and folded over 1,035 SMD frames. This target has three domains: Domain 1 (res. 1–83 and 205–240), Domain 2 (res. 84–204), and Domain 3 (res. 241–254). Five SMD trajectories of this protein have two different folding pathways (Fig. 4B). Four of the five trajectories (blue, red, purple, and orange) followed highly similar folding pathways, whereas the remaining trajectory exhibited a slightly different order of domain formation. In the dominant folding pathway observed in four of the five trajectories, folding begins with the formation of the first portion of Domain 2 (res. 84–170) between SMD frames 226 and 528. This is followed by the formation of a segment of Domain 1 (res. 20–65) between frames 530 and 571. The remaining portion of Domain 2 then folds between frames 580 and 728, after which Domain 1 reaches its native conformation between frames 730 and 979. Finally, Domain 3 forms during the late stage of folding, between frames 980 and 1035. In the alternative pathway (green) observed in the remaining trajectory, Domain 2 folds almost completely between SMD frames 190 and 620. Subsequently, the two segments of Domain 1 folded concurrently between frames 630 and 979, followed by the formation of Domain 3 between frames 980 and 1035. Despite this variation in the order of intermediate folding events, all trajectories exhibit the same overall trend: Domain 2 forms first, whereas Domain 3 forms last. This consistency suggests that the relative ordering of the earliest and latest folding events is robust across trajectories.

In the predicted pathway by PathFold, formation of Domain 2 was identified at frame 4, corresponding to SMD frame 593 with a structural similarity of 0.795. The subsequent formation of the Domain 1 segment (residues 20–65) was captured at PathFold frame 5, corresponding to SMD frame 778 with a similarity of 0.863. Finally, the formation of Domain 3 was identified at frame 9, which closely matches SMD frame 1012 with a high structural similarity of 0.963. Overall, PathFold accurately reproduced the hierarchical folding process of 5UJD_B, correctly identifying the early formation of Domain 2 and the late formation of Domain 3.

The third example is the extracellular solute-binding protein (PDB ID: 5SV6_A) ^45^, which contains 246 residues and an SMD trajectory of 995 frames (**Fig. 3C**). This protein consists of two domains: Domain 1 (D1; res. 1–77 and 195–246) and Domain 2 (D2; res. 88–194), connected by a flexible β-sheet linker (residues 78–87). Analysis of the five SMD trajectories revealed three folding variations (**Fig. 4C**). The five MD trajectories differ primarily in the folding behavior of the D1 subregion spanning residues 6–60 (Fig. 4C). MD1 (blue) and MD5 (orange) follow nearly identical folding pathways. In both trajectories, D2 folds first, followed by the β-sheet linker. Within D1, the 6–60 subregion is the final region to fold (SMD frames 788–808 for MD1 and 807–875 for MD5).The other three trajectories deviate from this pathway. In MD2 (red) and MD4 (purple), the 6–60 D1 subregion forms as an independent folding intermediate much earlier, at SMD frames 250–403 and 711–766, respectively, before the remainder of D1 is completed. In contrast, in MD3 (green), the 6–60 D1 subregion never forms a distinct intermediate but instead fold progressively together with the remainder of D1 between SMD frames 789 and 876.

The predicted trajectory by PathFold most closely resembles the alternative folding pathway observed in MD2 (red) and MD4 (purple). It first predicts partial folding of D2 at frames 3–4. Next, the D1 6-60 subregion forms as a distinct folding unit at frame 6, matching SMD frame 713 with a contact similarity of 0.822. Finally, the β-sheet linker and the remaining portions of D1 fold during frame 7–9.

Finally, we present an example where the predicted folding path by PathFold somewhat differs from the SMD trajectories (**Fig. 3D**, **Fig. 4D**). The target is the *Arabidopsis thaliana* Toc75 polypeptide transport-associated (POTRA) domains (PDB ID: 5UAY_A) ^46^, which consists of 302 residues and a 1,230-frame SMD trajectory. The structure contains three tandem POTRA domains: POTRA1 (res. 34–103), POTRA2 (res. 104–193), and POTRA3 (res. 194–302). The five SMD simulations were highly consistent with an average trajectory similarity score of 0.937 (**Fig. 4D**). The SMD trajectories show a sequential domain folding process. POTRA2 folds first between SMD frames 430–543, followed by POTRA3 between frames 728–858. Then, POTRA1 folds last between frames 879–1074, completing the native three-domain architecture.

On the other hand, PathFold predicted that a part of POTRA3 folded first (frame 5), followed by POTRA2. POTRA1 was folded last, which is consistent with SMD. At frame 4, PathFold predicted a partially folded intermediate centered on residues 151–193 of POTRA2. However, the corresponding stage of the SMD trajectory contains a rapidly folding POTRA2 and does not exhibit a comparable intermediate. At frame 5, PathFold predicted POTRA3 to be largely folded, whereas the corresponding stage of the SMD trajectory is dominated by the rapid folding of POTRA2. POTRA2 subsequently folds at frame 6. Finally, frame 7–9 capture the gradual folding of POTRA1, which agrees well with the SMD trajectories, yielding contact similarities of 0.940 and 0.958 with SMD frames 1113 and 1164, respectively.

### Folding Pathway Prediction by three models of PathFold

Here, we compare the overall alignment scores of three PathFold models that use the previous one, three, or six intermediate frames as input to predict the next structure along the folding pathway (PathFold-1, PathFold-3, and PathFold-6, respectively) (**Fig. 5**; individual data provided in **Supplementary Tables S4–S6**). The first panel (**Fig. 5A**) shows the distributions of alignment scores and reordered alignment scores. For the reordered alignment scores, the predicted trajectory is reordered according to the similarity of each intermediate structure to the AF2-predicted folded structure, and the alignment scores are recomputed using PathScorer. This reordering is feasible during inference because the AF2 structure is already generated to provide the input representation. Among the three models, PathFold-3 showed overall the highest average alignment score, followed by PathFold-1 and PathFold-6. The average alignment scores of the three models were 0.815, 0.804, and 0.733, for PathFold-3, −1, and –6, respectively. When the predicted intermediate structures were reordered according to their similarity to the native structure, the alignment scores increased for all three models. The average reordered alignment scores were 0.840, 0.848, and 0.848 for PathFold-1, PathFold-3, and PathFold-6, respectively. The largest improvement was observed for PathFold-6, indicating that this model more frequently generates trajectories with non-monotonic folding progression.

**Figure 5.**
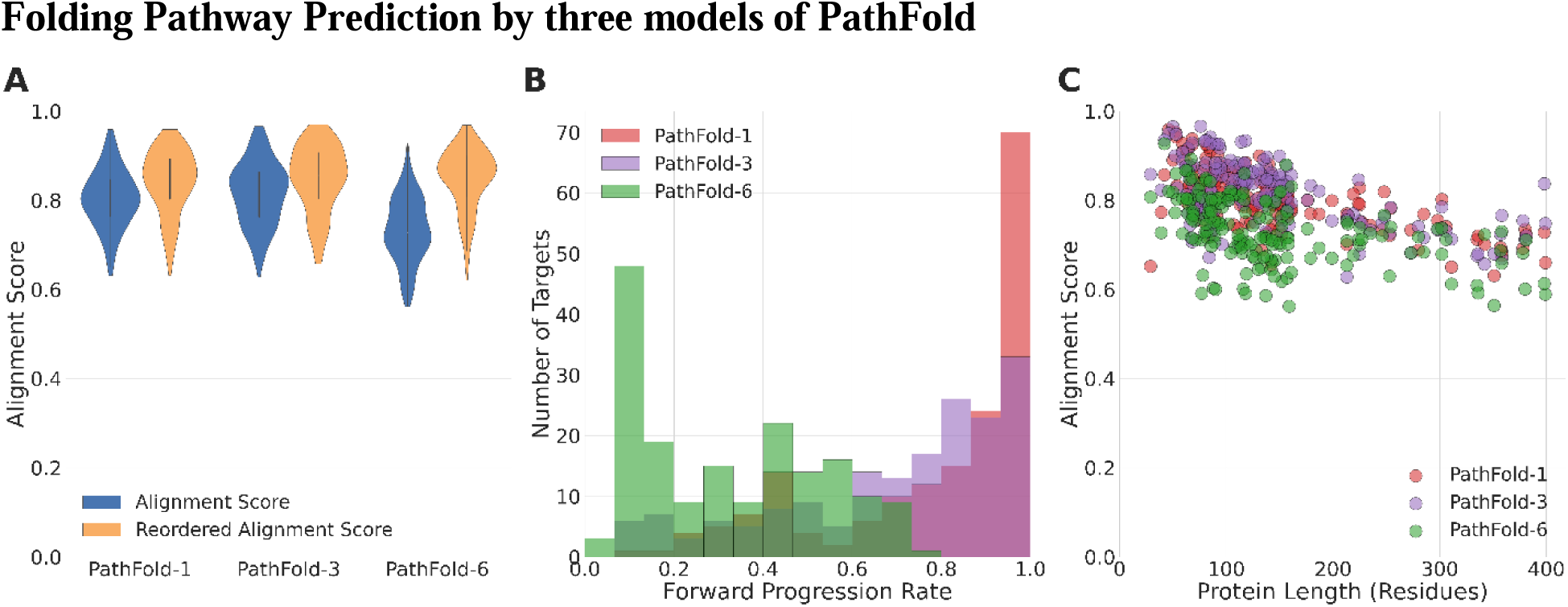
Alignment scores of predicted pathways by PathFold. Pathway prediction performance of three PathFold models using one, three, or six previous frames as input (PathFold-1, PathFold-3, and PathFold-6, respectively) was evaluated on 175 test proteins. **A**, Violin plots of alignment scores and reordered alignment scores. Reordered alignment scores were computed after reordering the predicted trajectories based on the contact-map similarity of each intermediate structure to the folded state (see text). Embedded box plots indicate the median, interquartile range, and 1.5 × interquartile range. Mean alignment scores were 0.804, 0.815, and 0.733 for PathFold-1, PathFold-3, and PathFold-6, respectively; the corresponding reordered alignment scores were 0.840, 0.848, and 0.848. **B**, Distribution of forward progression rate for PathFold-1 (red), PathFold-3 (purple), and PathFold-6 (green). The forward progression rate is the proportion of the 10 candidate structures generated at each iteration that exhibit forward folding progression (see text). **C**, Alignment scores as a function of protein length for PathFold-1 (red), PathFold-3 (purple), and PathFold-6 (green).

This trend is more evident in **Fig. 5B**, which shows the forward progression rate, a measure of whether predicted structures move closer to the native structure during each prediction step. At each iteration, the forward progression rate is defined as the fraction of the 10 candidate structures that are closer to the native structure than the input frame(s). This value is then averaged over all iterations in a trajectory. Closeness to the native structure is measured by the Dice similarity between the contact maps of the predicted and native structures. PathFold-6 exhibited a distribution shifted toward lower forward progression rates than the other two models. The average forward progression rates were 0.784, 0.678, and 0.325 for PathFold-1, PathFold-3, and PathFold-6, respectively. A prediction step was considered to move backward when all 10 candidate structures generated at an iteration were farther from the native structure than the input frame. This occurred in 28.9% of iteration steps for PathFold-6, which is substantially more frequent than PathFold-1 (15.1%) and PathFold-3 (13.0%). We also found that trajectories predicted by PathFold-6 more frequently exhibited non-native-like folding behavior, such as excessively rapid or slow conformational changes along the pathway (**Supplementary Table S7**). To quantify these deviations, we implemented the native-like trajectory score in PathScorer, which penalizes the three types of non-native-like behavior described above (Eq. 5 in Methods). Using this metric, trajectories generated by PathFold-6 received substantially lower scores than those generated by PathFold-1 and PathFold-3 (**Supplementary Figure S8**).

Interestingly, PathFold-6 performed the worst, even though one might expect it to perform best because access to a longer history of previous frames should provide richer information about the folding trajectory. A likely explanation is the reduced amount of effective training information available to PathFold-6. Because each prediction conditions on the six preceding intermediate states, many training trajectories—particularly those of short proteins—do not contain sufficient frames to provide all six inputs, requiring the missing inputs to be padded with zeros. Consequently, 38.7% of the training samples for PathFold-6 contain one or more padded inputs, compared with 19.4% for PathFold-3 and only 6.5% for PathFold-1. The frequent use of padded inputs likely reduced the amount of informative context available during training and contributed to the poorer performance of PathFold-6.

**Fig. 5C** shows the alignment score as a function of protein length. The score decreases slightly with increasing protein length. This trend is expected because longer proteins have more elements to fold, which often contain multiple domains.

### Φ-value Analysis for Folding Trajectories

Finally, we examined PathFold-predicted folding trajectories for three proteins with experimentally measured Φ-values, a well-established measure for characterizing protein-folding transition states and identifying residues that form native-like interactions early in the folding process^12^. Experimentally, Φ-values are determined by point mutations and calculated as the ratio of the mutation-induced change in the activation free energy of folding to the corresponding change in overall protein stability. Values near 1 generally indicate that interactions involving the mutated residue are largely native-like in the transition-state ensemble, whereas values near 0 indicate that they remain largely denatured-like.

To compare PathFold trajectories with experimental Φ-values, we computed a residue-level folding score ranging from 0 to 1 based on the timing of native-contact formation (see Methods). A higher score indicates that a residue establishes its native-like local environment earlier in the predicted folding trajectory.

We analyzed three well-characterized proteins with experimental Φ-values available for more than a dozen residues: the FBP28 WW domain (PDB ID: 1E0L)^36^, the cold shock protein B (CspB; PDB ID: 1CSP)^37^, and ubiquitin (PDB ID: 1UBQ)^38^. Experimental Φ-values were obtained from works Petrovich et al.^47^, Garcia-Mira et al.^48^, Went and Jackson.^49^, following the procedure used by Ooka et al.^50^. The data are provided in **Supplementary Tables S8–S10**.

The first example (Fig. 6A), the FBP28 WW domain (PDB ID: 1E0L) is a 37-residue, three-stranded β-sheet protein. Experimentally, Gly16 (red) and Thr18 (yellow) have high Φ-values of 1.00 and 0.84, respectively, indicating early formation of the first β-hairpin and the beginning of β-strand 2 during folding^47^. PathFold qualitatively reproduced this pattern. At the timestamp 1, the highest contact fractions were concentrated around β-turn I, including Ala14 (residue fraction score: 0.50) and Asp15 (0.67), while contacts within β-strands 1 and 2 remained largely unformed (0.08 to 0.17). By timestamp 2, contact formation had expanded throughout the first β-hairpin. Consistently, the PathFold folding-score profile (right top panel) peaked around β-turn I, with high scores for Ala14 (0.96), Asp15 (1.00), Gly16 (0.94), and Thr18 (0.66), which shows a similar profile as the Φ-value profile. Overall, residues 8–23 in β-strand 1 and 2 had a substantially higher mean folding score (0.79) than the remaining residues (0.43), supporting qualitative agreement between the PathFold trajectory and the experimentally identified high-Φ region.

**Figure 6.**
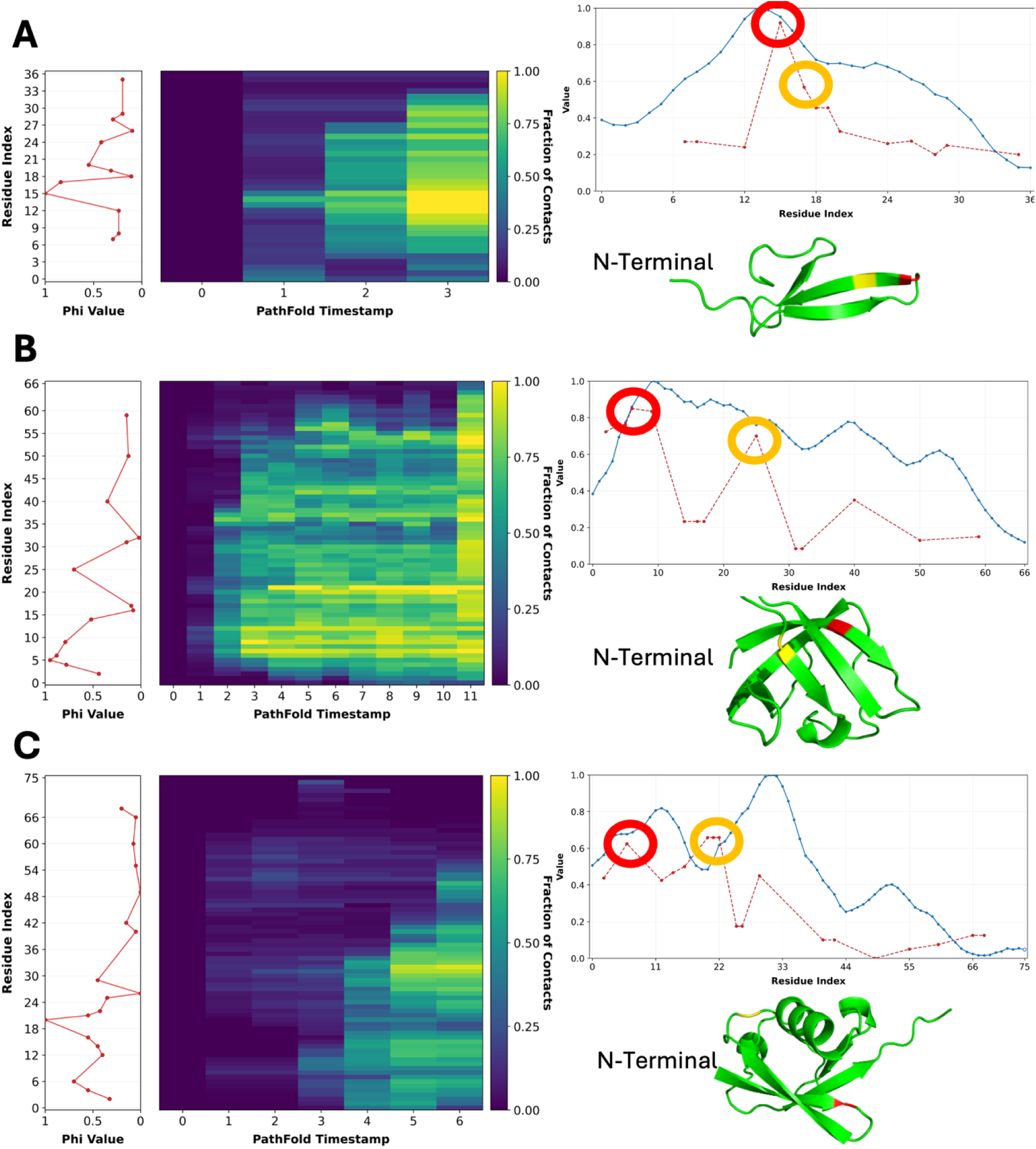
Comparison with experimental Φ-values. Folding events observed in PathFold-predicted trajectories were compared with experimental Φ-values for three proteins. Left: Experimentally measured Φ-values by residue; residues without experimental measurements are left unassigned. Middle: Heat map showing, at each PathFold time point, the fraction of contacts formed by each residue relative to its contacts in the AF2-predicted folded structure. Residue positions are shown on the y-axis, with the color scale ranging from 0 (dark purple) to 1 (yellow). Right, top: Comparison of the PathFold folding score (blue; see text) with experimental Φ-values (red dashed line). PathFold scores were smoothed along the sequence and normalized to 0–1, whereas experimental Φ-values were smoothed without normalization. Moving-average windows of two residues were used for the FBP28 WW domain and ubiquitin, and three residues for the cold shock protein. Right, bottom: AF2-predicted folded structure. **A**, FBP28WW domain (PDB ID: 1E0L, 37 residues). Red, Gly16; yellow, Thr18. **B**, cold shock protein (PDB ID: 1CSP, 67 residues). Red, Lys7; yellow, Val26. **C**, ubiquitin (PDB ID: 1UBQ, 76 residues). Red, Thr7; yellow, Asp21.

The second example is cold shock protein B (CspB; PDB ID: 1CSP), a 67-residue, five-stranded β-barrel (Fig. 6B). Experimental Φ-values indicate a strongly polarized transition state, with high values concentrated in the N-terminal β1–β3 sheet, particularly at Lys5, Val6, Lys7 (red), Asn10, and Val26 (yellow), but not in the C-terminal β4–β5 sheet^48^. The Lys7–Asp25 interaction has also been implicated in early docking of β3^51^. PathFold qualitatively reproduced this pattern. At timestamp 2, β1–β3 showed a higher mean contact fraction than the remaining regions (0.38 vs. 0.22), and the Lys7–Asp25 contact was formed. The folding-score profile similarly favored β1–β3 (β1: residue 2–10, β2: 14–19, and β3: 26–30), with a mean score of 0.72 compared with 0.55 for the remaining residues. Particularly high scores were observed for Lys7 (0.87), Asn10 (0.98), Asp25 (0.73), and Val26 (0.68), supporting early formation of the N-terminal sheet consistent with the experimental Φ-values.

The third example is ubiquitin (PDB ID: 1UBQ), a 76-residue α/β protein (Fig. 6C). Experimental Φ-values indicate a polarized transition state involving the N-terminal β-hairpin (residues 1–17) and major α-helix (residues 23–34), with high Φ-values at Thr7 (red) and particularly Asp21 (yellow), whereas residues 41–69 showed uniformly low values^49,52,53^. PathFold reproduced this overall N-terminal-first folding pattern, although residue-level agreement was less pronounced than in the previous examples. By timestamp 4, the β-hairpin and α-helix reached mean contact fractions of 0.45 and 0.43, respectively, forming a native-like arrangement. Consistently, residues 1–35 had a substantially higher mean folding score than residues 36–76 (0.66 vs. 0.23). Although scores dipped around residues 18–23 due to their extensive contacts with the C-terminal region, residues 31–35 showed particularly high scores (mean 0.91). Overall, PathFold predicted early assembly of the N-terminal β-hairpin–helix region, qualitatively consistent with the experimentally inferred folding nucleus.

## Discussion

We presented PathFold, a diffusion-based framework for predicting protein folding trajectories from an unfolded state to the native conformation using only sequence information. To our knowledge, this is the first deep learning method designed to generate complete, ordered protein folding pathways. PathFold produces progressive intermediate conformations and trajectories that show substantial agreement with SMD simulations. Furthermore, residue-level folding scores derived from the predicted trajectories generally reproduce trends in experimental Φ-values, suggesting that the generated pathways capture meaningful features of the folding process.

A fundamental challenge in developing data-driven models of protein folding is the lack of large-scale atomistic folding trajectory datasets. Complete folding processes are difficult to observe experimentally at atomic resolution, while generating spontaneous folding trajectories computationally at sufficient scale remains prohibitively expensive. We therefore used reversed SMD unfolding trajectories as a practical surrogate for developing and evaluating trajectory-learning methods. Although these trajectories are not equivalent to genuine folding pathways and do not reproduce their kinetics or thermodynamics, they provide ordered conformational transitions for establishing whether deep learning can learn structural trajectories rather than simply their endpoints. PathFold thus provides a methodological foundation that can be applied to genuine folding trajectories as suitable datasets become available.

We also developed trajectory alignment and similarity metrics to quantitatively compare molecular pathways, where similar structural events may occur at different time points. These metrics align trajectories by structural progression rather than simulation time, enabling comparison of intermediate conformations and folding events. Together, PathFold and these evaluation methods provide a framework for data-driven modeling of protein folding and other dynamic molecular processes.

Several directions remain for future work. First, PathFold currently operates at the Cα level and therefore does not explicitly model side-chain interactions that influence folding dynamics. In addition, although the current folding progression criterion stabilizes trajectory generation and prevents abrupt structural transitions, incorporating more physically grounded energy or thermodynamic constraints could further improve the realism and robustness of the predicted pathways. Finally, the framework is readily extendable to other types of molecular trajectories, such as protein–ligand binding and conformational transitions, provided suitable training datasets are available.

More broadly, machine learning approaches such as PathFold highlight the growing potential of AI-driven methods to complement traditional biophysical simulations. Recent advances in ML-based molecular modeling have demonstrated the ability to accelerate conformational sampling^54^, and learn generalized molecular dynamics patterns^55^ that remain computationally challenging for conventional MD simulations alone. Such developments may ultimately contribute to large-scale simulations of cellular systems^56^, enabling more efficient studies of folding, binding, conformational transitions, and biomolecular assembly dynamics.

## Acknowledgements

We thank Prof. Steven Hayward for helpful discussions and technical advice on protein trajectory morphing. This work was partly supported by the National Institutes of Health (R35GM158267, R01GM133840, R21AI187928) and the National Science Foundation (IIS2211598, DMS2151678, DBI2146026 and DBI2422620).

## Author contributions

DK conceived the study. Z.Z designed and implemented the PathFold pipeline. Y.K. prepared SMD data. Z.Z performed the computation and analyzed the data. N.I designed and implemented alignment scoring. Z.Z and Z.X analyzed Φ-values experiments. P.P participated in analyzing folding trajectory data. Z.Z., Y.K., N.I, Z.X. drafted the paper. D.K. critically edited it. All the authors read and approved the manuscript.

## Methods

### Dataset Construction

We used MD simulations as described in our previous work ^40^. The dataset was curated from the January 2019 PISCES ^57^ release, restricted to proteins with <25% sequence identity and resolution ≤2.5 Å. Chains shorter than 25 residues, those containing non-standard residues, and structures released after May 14, 2018 (CASP13 cutoff) were excluded. Structures with gaps >2 residues were removed; smaller gaps were modeled using MODELLER^58^, retaining only those with Cα-RMSD <2.0 Å relative to the original structures. Knotted proteins and entries failing during pulling simulations were further excluded, yielding 6,718 protein chains. For computational efficiency, 1,290 proteins with ≤400 residues were randomly selected for training, validation, and testing.

### Steered Molecular Dynamics (SMD) Simulation

The SMD simulation protocol was based on our previous work^40^. All molecular dynamics (MD) simulations were performed using NAMD^59^ with a 1 fs timestep. Each protein was first parameterized using the CHARMM36 force field^60^ via the autopsf tool in VMD^61^, with disulfide bond formation disabled. For SMD setup, the Cα atom of the N-terminal residue was fixed and the Cα atom of the C-terminal residue was pulled along the normalized vector connecting the two termini. Systems were energy-minimized for 10,000 conjugate gradient steps using a Generalized Born implicit solvent (GBIS) model. Production SMD simulations were then performed with a spring constant of 7 kcal/mol/Å² and a constant pulling velocity of 0.001 Å/fs applied to the C-terminal Cα. The total simulation length was determined per protein as

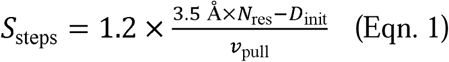

where *Nres* is the number of residues, *D*_init_ is the initial Cα–Cα distance, and the factor 1.2 provides a 20% safety margin assuming 3.5 Å per residue in an extended chain. For each protein, the entire protocol, consisting of energy minimization followed by SMD, was independently repeated five times starting from the same initial PDB structure. Coordinates were saved every 1 ps for analysis.

### Trajectory alignment score

To quantify the similarity of two folding trajectories, we developed PathScorer, which performs global alignment of two trajectories using dynamic programming (DP) ^62^. At each alignment step, structural similarity is evaluated using contact map–based scores between conformations. While inspired by previous work that applied DP to compare MD trajectories^63^, PathScorer is specifically designed for comparing complete folding trajectories spanning unfolded to native conformations. Instead of global root mean square deviation (RMSD), it uses normalized contact map similarity and trajectory-aware gap penalties, enabling robust alignment of folding pathways that differ in rate, duration, and local directionality.

Formally, let a MD trajectory be *T* = {*t*_1_ *, t*_2_ *, …, t_n_*}, captured at intervals of *dt*, where *t_i_* represents the protein structure at the *i*^th^ time point. Similarly, the predicted trajectory by PathFold is *P* = {*p*_1_ *, p*_2_ *, …, p_m_*}, but generated at a step size of Δ*t*. We aim to compute a score *score(P, T)* that quantifies the similarity between the two trajectories. In our method, *dt < Δt*, which implies *n > m*. To compare the trajectories, we sample subsequences of equal length *k*(=*min*(*n,m*) *= m*) from each: 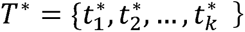 and 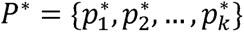. Using a similarity metric 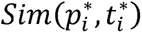, we compute the similarity between matched frames, potentially including penalty terms to encourage selection of a reasonable and meaningful subsequence.

To compare two conformations, a root mean square deviation (RMSD) is not well suited for comparing folding trajectories because large conformational changes can produce highly variable values that do not reliably reflect folding progress. Instead, we use contact map similarity, which naturally captures the formation of residue–residue interactions during folding. Two residues were considered in contact if their Cα atoms were within 14 Å, excluding pairs separated by fewer than three sequence positions. Similarity between two structures was defined as the number of shared contacts, normalized by the contacts common to the native folded structure:

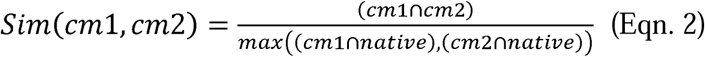

where cm1, cm2 are two contact maps from two different folding trajectories, and native corresponds to the final folded structure. ∩ takes the common contacts between them. When counting shared contacts, we only consider the residue-pairs in contact and ignore the ones that are not (e.g. stretched conformation). Normalizing by the shared contact of the folded native structure, makes the score upper bounded by 1 for perfect matches.

Because predicted and ground-truth trajectories contain different numbers of frames, alignments invariably include gaps. Because PathFold predicts structures at 50-frame intervals, no penalty was applied to gaps spanning ≤50 frames. For longer gaps, the extension penalty was defined by the structural distance across the gap rather than by a conventional linear model, which is ill suited to the highly variable rates of conformational change observed along folding trajectories. Structural distance was quantified using the Dice distance between contact maps, which empirically provided a robust measure of conformational change.

The recurrence relations for the DP are as follows:

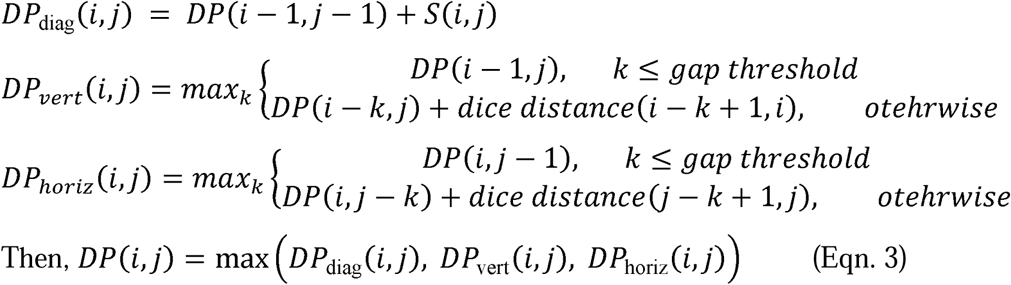

where the gap threshold is set to 50. Using this dynamic programming recurrence, the global alignment score *DP*(*n,m*) is computed and normalized by the number of matches, i.e., the length of the shorter trajectory, to yield a score between 0 and 1. Therefore,

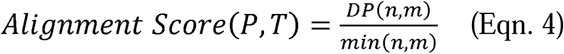

In addition, we developed a new alignment metric, the native-like trajectory score, which penalizes non-native-like behavior in predicted folding trajectories. This score is used only to evaluate pathway alignments and does not affect the alignment procedure itself. Although the alignment method described above provides a robust measure of trajectory similarity, PathFold occasionally generates trajectories that progress too rapidly, too slowly, or even reverse their folding progression. The native-like trajectory score introduces heuristic penalties for these behaviors based on changes in similarity to the native folded structure between consecutive predicted frames. Specifically, changes exceeding a predefined threshold are classified as rapid transitions, changes below a threshold as slow transitions, and negative changes as reversals in folding progression. The corresponding penalties were defined as follows:

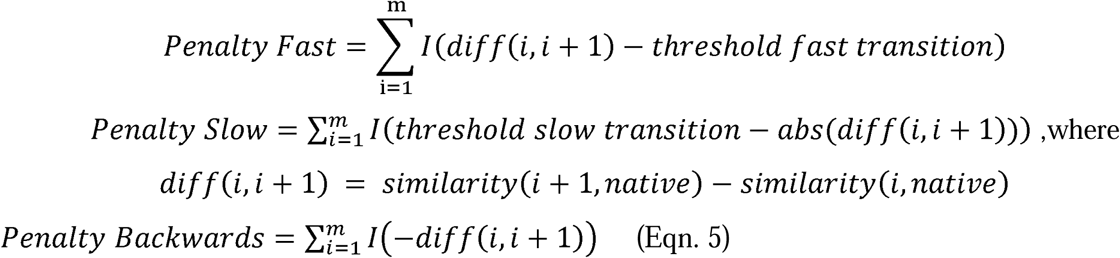

The threshold values for rapid and slow transitions were set to 0.2 and 0.01, respectively. These values were determined heuristically based on a manual review of ∼30 predicted trajectories. Specifically, transitions with a similarity increase greater than 20% between consecutive frames were classified as rapid, whereas those with an increase of less than 1% were classified as slow and penalized.

The native-like trajectory score is obtained by combining the DP alignment score with these additional penalties:

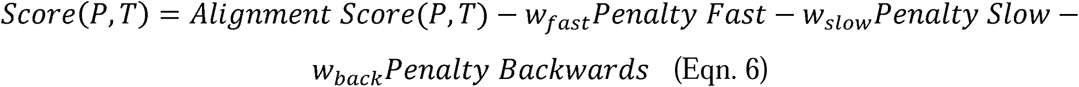

We used *w_fast_, w_Slow_, w_back_* as 0.1. These weights were likewise selected based on a heuristic evaluation. We manually reviewed poorly predicted trajectories and assigned weight values designed to distinctively separate their score distribution from that of ideal predictions.

### Training PathFold model

PathFold is a denoising diffusion probabilistic models (DDPMs) framework conditioned on AF2 embeddings and an intermediate folding state. PathFold generates the next intermediate folding states towards folding the protein.

The input to PathFold consists of a previous state and a target state, represented as Cα structures extracted from SMD simulations that are separated by 50 frames. During training, PathFold is optimized to predict forward folding progression, where the less folded structure serves as the previous state and the more folded structure serves as the target state. The Cα coordinates are first converted into Frenet–Serret (FS) frames, which represent each residue using a rotation matrix and translation vector derived from the backbone geometry. These frames are processed by an SE(3)-invariant encoder, which produces single and pair encodings based on per-residue frames, sinusoidal encodings of residue indices, and the diffusion timestep. The resulting single and pair encodings are then fused with AF2 single and pair representations. Specifically, given single embeddings of size *L* × *H* and pair embeddings of size *L* × *L* × *H*, where *L* is the protein length and *H* is the hidden dimension, linear projection layers are applied to map the AF2 representations into the same embedding dimension as the invariant encoder outputs. To encode the intermediate folding state, the previous structure is converted into a contact map, where two residues are considered in contact if their Cα–Cα distance is less than 10 Å. The contact map is processed using two ResNet blocks composed of 3×3 convolutional layers, producing pair embeddings of size *L* × *L* × *H*. These embeddings are then combined with the projected AlphaFold pair representations through element-wise addition, providing structural context for the decoder.

PathFold was implemented in PyTorch. Training required approximately eight weeks on a single NVIDIA RTX 6000 GPU with 48 GB of memory. Our implementation builds upon the Genie framework^39^, whose architecture we adapted for folding trajectory prediction. We trained the model using the AdamW optimizer with a learning rate of *1e10*^—3^, a batch size of 4, and gradient accumulation over 8 steps. AF2 was run using the official implementation available at https://github.com/google-deepmind/alphafold to generate single and pair representations used as structural priors for PathFold.

### Folding Trajectory Generation

At each iteration, PathFold simultaneously generates ten candidate structures in a single diffusion sampling pass. To promote smooth and physically plausible folding pathways, the candidates are evaluated using with the Dice similarity to the folded structure, which is defined as:

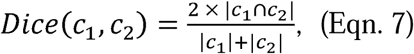

where *c*_1_ and *c*_2_ are the contact map of candidate structure and folded structure, respectively. Two residues were considered in contact if their Cα atoms were within 14 Å. A candidate is accepted only if its similarity to the folded structure predicted from the AF embeddings increases relative to the previous iteration and the increase remains below a predefined threshold (0.01), preventing large structural jumps. Among all accepted candidates, the structure showing the greatest forward progression toward the folded state is selected. If no candidate satisfies these criteria, a fallback strategy chooses the structure with the smallest decrease in similarity. The selected structure is then used as the starting point for the next iteration, and the process repeats until a similarity threshold of 0.95 to the folded conformation is reached. This procedure encourages monotonic and gradual folding progression while preserving meaningful intermediate states.

### Analyzing Folding Trajectories at a Residue Level

From a predicted folding trajectory by PathFold, we evaluated the nativeness of residues along the trajectory by considering the fraction of native contacts formed at that time. For residue *i* in a trajectory frame *j*, the fraction of native contacts is computed as

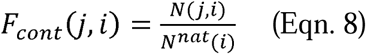

where *N^nat^*(*i*) is the number of contacts formed by residue *i* in the native structure (AF2-predicted structure) and *N(j, i)* is the number of those contacts that are presented at frame *j.* Consistent with Eqn. 2, two residues are considered in contact if their Cα-Cα distance is below 14 Å. Residue pairs are not considered if they are within three residues on the sequence. The residue folding score of a residue *i* is then computed by accumulating F_cont_ over all the frames:

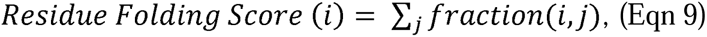

A residue folding score will be close to 1 if it establishes its native contact environment in an early stage and retains it in the folding trajectory.

